# A Prediction Tool for Plaque Progression Based on Patient-specific Multi-Physical Modeling

**DOI:** 10.1101/2020.09.17.301218

**Authors:** Jichao Pan, Yan Cai, Liang Wang, Akiko Maehara, Gary S. Mintz, Dalin Tang, Zhiyong Li

**Affiliations:** School of Biological Sciences and Medical Engineering, Southeast University, Nanjing Jiangsu, China; The Cardiovascular Research Foundation, New York, New York, USA; Mathematical Sciences Department, Worcester Polytechnic Institute, Worcester, MA 01609, USA; School of Mechanical, Medical and Process Engineering, Queensland University of Technology, Brisbane, QLD 4001, Australia; Wolfson College, University of Cambridge, Barton Road, Cambridge CB3 9BB, UK

## Abstract

Atherosclerotic plaque rupture is responsible for a majority of acute vascular syndromes and this study aims to develop a prediction tool for plaque progression and rupture. Based on the follow-up coronary intravascular ultrasound imaging data, we performed patient-specific multi-physical modeling study on four patients to obtain the evolutional processes of the microenvironment during plaque progression. Four main pathophysiological processes, i.e., lipid deposition, inflammatory response, migration and proliferation of smooth muscle cells (SMCs), and neovascularization were coupled based on the interactions demonstrated by experimental and clinical observations. A scoring table integrating the dynamic microenvironmental indicators with the classical risk index was proposed to differentiate their progression to stable and unstable plaques. The heterogeneity of plaque microenvironment for each patient was demonstrated by the growth curves of the main microenvironmental factors. The possible plaque developments were predicted by incorporating the systematic index with microenvironmental indicators. Five microenvironmental factors (LDL, ox-LDL, MCP-1, SMC, and foam cell) showed significant differences between stable and unstable group (p < 0.01). The inflammatory microenvironments (monocyte and macrophage) had negative correlations with the necrotic core (NC) expansion in the stable group, while very strong positive correlations in unstable group. The inflammatory microenvironment is strongly correlated to the NC expansion in unstable plaques, suggesting that the inflammatory factors may play an important role in the formation of a vulnerable plaque. This prediction tool will improve our understanding of the mechanism of plaque progression and provide a new strategy for early detection and prediction of high-risk plaques.

**Author summary:** Besides the traditional systematic factors, the influences of the local microenvironmental factors on atherosclerotic plaque progression have been demonstrated. Mathematical and computational modeling is an important tool to investigate the complex interplay between plaque progression and the microenvironment, and provides a potential way toward the prediction of plaque vulnerability according to the comprehensive evaluation of both morphological and/or biochemical factors in tissue level with microenvironmental factors in cellular level. We performed patient-specific multi-physical modeling study on four patients to obtain the evolutional processes of the microenvironment during plaque progression and predicted the possible plaque developments. A scoring table integrating the dynamic microenvironmental indicators with the classical risk index was proposed to differentiate their progression to stable and unstable plaques. Based on patient-specific imaging data, the mathematical model will provide a novel method to predict the changes of plaque microenvironment and improve ability to access the personal therapeutic strategy for atherosclerotic plaque.

## Introduction

Atherosclerosis is the process in which plaques, consisting of lipids, monocytes/ macrophages (MΦs), vascular smooth muscle cells (SMCs) and calcium, are built up in the walls of arteries as a chronic inflammatory response. Coronary atherosclerosis can cause myocardial infarction and heart failure [1]. Although multiple systemic atherosclerotic risk factors have been identified, including age, family history, hypertension, hypercholesterolemia, diabetes and so on, advances in molecular and cellular researches have significantly enhanced our understanding of the influence of the local biochemical/biophysical microenvironment on atherosclerotic plaque progression. It is now acknowledged that atherogenesis is a function of both systemic atherosclerotic risk factors and the local microenvironmental stimuli.

Multiple microenvironmental factors contribute to plaque formation and progression, including arterial mechanics, hemodynamics, matrix composition, lipid deposition, inflammation and neovascularization from vasa vasorum (VV) [2–7]. The cellular components (MΦs, SMCs, endothelial cells, etc.) and their extracellular environment exist in a state of dynamic reciprocity, which influences all stages of plaque progression and determines the plaque fate as to whether it would develop to a vulnerable phenotype. For example, in the early phase, hypercholesterolemia conditions increase low-density lipoprotein (LDL) infiltration and retention into the injured endothelial layer in response to disturbed blood flow pattern, leading to the accumulation of inflammatory cells by the release of pro-inflammatory factors, such as monocyte chemoattractant protein (MCP-1) [6,8,9]. Meanwhile, vascular SMCs undergo phenotypic dedifferentiation that from quiescent phenotype to a synthetic and activated phenotype, in response to pro-inflammatory cytokines and oxidized LDL (ox-LDL) [10–12]. In addition, SMCs express a number of pro-inflammatory chemokines in turn and even phenotypically transform to macrophage-like cells [10,11]. Another important pathological process involved in the coupled interactions among the microenvironmental components within the plaque is intraplaque angiogenesis [2], which refers to the disorganized and abnormal neovascularization from adventitial VV in response to the hypoxic microenvironment in the thickening intimal. Intraplaque angiogenesis together with the associated intraplaque hemorrhage (IPH) contributes to the accumulation of inflammatory cells and lipoproteins through the leaky angiogenic microvessels, which are vital characteristics of atheroma microenvironment in vulnerable plaques [2,5]. To summarize, an understanding of plaque development and progression requires the elucidation of multiple factors and their interactions within the microenvironment.

Unfortunately, there are fundamental gaps in our knowledge of the underlying mechanisms that contribute to the influences of plaque microenvironmental factors on atherogenesis in vivo. The main reason for this is that the microenvironment that determines the disease activity, including lipid deposition, inflammation, neovasculature and hemorrhage, cannot be assessed by non-invasive plaque-imaging techniques (such as ultrasound, CT and MR) which are widely used to obtain plaque morphological characteristics in clinic. Although advanced imaging techniques, including intravascular ultrasound (IVUS), optical coherence tomography (OCT) and PET/CT, now have permitted more accurate assessment of the plaque composition and even the atherosclerotic disease activity, the limited spatial resolution and lack of dynamic changes of microenvironmental factors give rise to barriers to quantitatively analyze the functional activity in atherosclerotic microenvironments.

Besides the developments of novel imaging techniques, mathematical modeling and numerical simulation have been proven useful for quantitatively assessing the dynamic changes of cellular and acellular components involved in the plaque microenvironment and predicting plaque growth and possible rupture [13,14]. In our previous studies, we have developed a multi-physical mathematical model by coupling lipid deposition, inflammation, neovascularization and intraplaque hemorrhage, to investigate the pathophysiological responses of plaques to dynamic changes in the microenvironment [15,16]. For the first time, the promotion of intraplaque angiogenesis on the accumulation of ox-LDL and macrophages in the plaque lesion and its quantitative contribution to the induction and progression of plaque destabilization were demonstrated by using this multi-physical model. This study is an application of our previously developed model for identifying the spatial-temporal dynamics and progression in coronary atheroma microenvironment based on patient-specific virtual histology(VH)-IVUS imaging data. The microenvironmental components in this model consist of four main parts, i.e., the plaque morphology and composition, the lipid deposition, the inflammation and the intraplaque neovascularization. To assess the personalized plaque microenvironment, simulations on a single patient data set are performed and the predicted changes of plaque composition are validated with the follow-up data from the same patient at the later time points. This study aimed (a) to describe the dynamic quantitative changes of the main factors in plaque microenvironment; (b) to predict the plaque development according to both systemic risk factors and microenvironmental factors; and (c) to investigate the distinct role of lipid and inflammatory microenvironment in the formation of a vulnerable plaque.

## Results

### Dynamic variation of the microenvironmental factors

The distribution changes of the three main variables (ox-LDL, MΦs and IPH) in four patients (P1-P4) are displayed in Fig 1, which represent the critical contributors in microenvironment to atherogenesis. An animation of the dynamics of all the variables involved in the simulation can be found in the Supplementary Material. At the early stage of the simulation (T1), most ox-LDL concentrated near the inner intima, while the area of high ox-LDL concentration gradually enlarged in the thickening intima (follow-up, T2) due to the extravasation of lipoprotein from the neovasculature, resulting in the lipid deposition near the outer intima in the plaque (at the end of simulation, T3). Similar to the lipid deposition, the exacerbating inflammation were found during plaque progression. A high concentration of IPH was observed in the necrotic core (NC) area throughout the plaque development, especially, at the shoulder of NC in P2.

**Fig 1.**
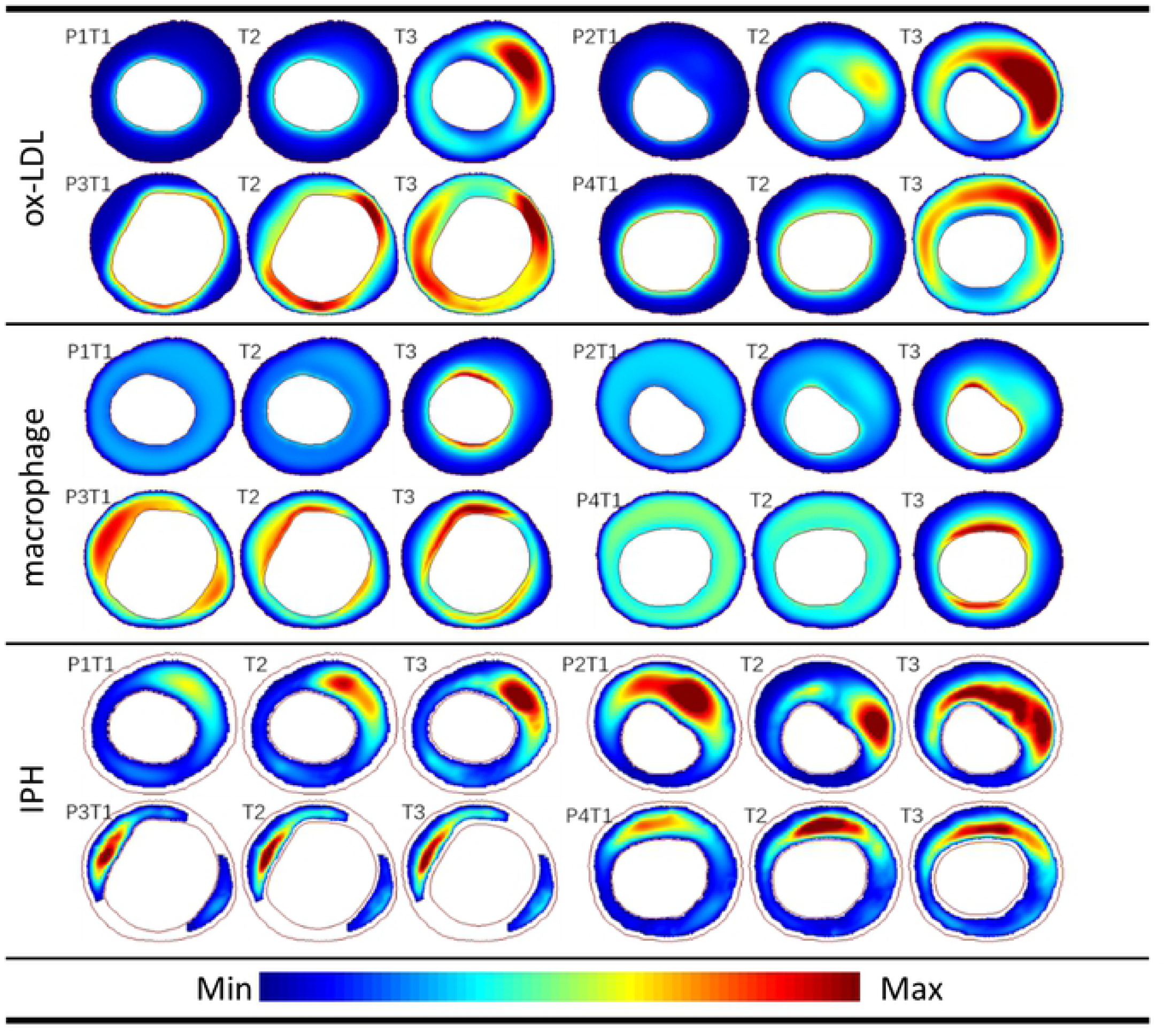
The progression of the main plaque microenvironmental factors. The values in all figures of each factor were normalized to the range of 0 to 1.

The heterogeneity of plaque microenvironment for each patient was observed apparently from the growth history curves of the average values of the eight main variables (Fig 2). Although there were no significant differences among the development trends between the four patients, the variation and the heterogeneity of plaque microenvironment between individuals were remarkable. For example, both a mild lipid deposition and a severe inflammation were found in P3 (yellow dotted line), resulting in an indistinctive apoptosis of SMCs. Nevertheless, an opposite situation happened in P2 (blue dotted line), where a high level of lipid and a low level of inflammation yielded a significant decrease in SMCs. In the following section, the evaluation of microenvironmental factors for each patient is given in the scoring scale table (Table 1).

**Table 1.** The multi-factorial scoring scale

|  | PB | EI | LDL | HDL | WSS | MΦ | SMC | ox-LDL | Total | Prediction |
| --- | --- | --- | --- | --- | --- | --- | --- | --- | --- | --- |
| P1 | 3 | 2 | 3 | 1 | 1 | 1 | 2 | 2 | 15 | stable plaque with macrocalcification |
| P2 | 3 | 3 | 3 | 3 | 1 | 1 | 3 | 3 | 20 | vulnerable plaque |
| P3 | 1 | 3 | 1 | 1 | 3 | 3 | 2 | 2 | 16 | stable plaque with low growth rate of NC |
| P4 | 2 | 2 | 3 | 3 | 3 | 3 | 2 | 2 | 20 | vulnerable plaque |

The prediction result is based on the calculation of the total score and the VH-IVUS images. EI = eccentric index, WSS = wall shear stress.

**Fig 2.**
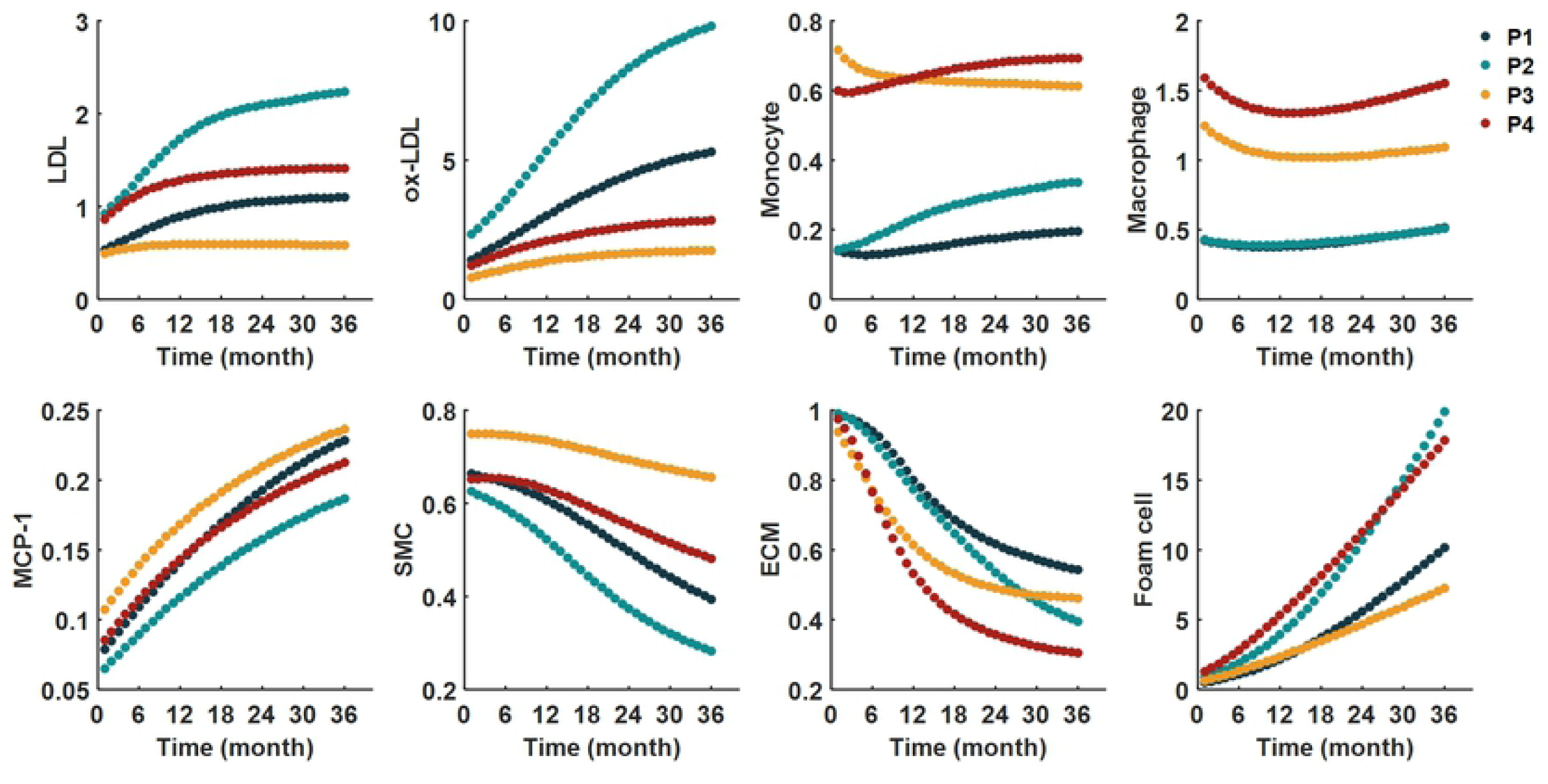
The concentrations of the plaque microenvironmental factors change with time. The values on the vertical axis are dimensionless.

### Prediction of plaque development

Fig 3A displays a comparison between the simulation results of the NC areas with the VH-IVUS images. The NC area at T1 was delineated based on the image segmentation of VH-IVUS data; while the expanded NC areas at T2 and T3 were determined according to the increasing areas of apoptotic SMCs (NC area was outlined with black/white border in the plaque). Different degrees of increase in plaque area and NC area were found in all four patients. Thin cap fibroatheroma (TCFA) with a large NC was observed especially in P2 and P4 at T3 stage, which suggests that these two plaques may be more likely to become vulnerable plaques. On the contrary, the flaky distributed calcification (white area shown in VH-IVUS) in P1 indicated a stable plaque. In addition, the quantitative curves of the growth rate of NC and plaque burden (PB) for the four patients are provided in Fig 3B. There was a remarkable increase in NC area of P4 at the early stage, while the NC areas in the other three patients increased steadily. Actually, the growth rate of NC in P4 was as high as 1.75 during the follow-up period according to the VH-IVUS images (between T1 and T2). According to the simulation results, the NC area in P4 increased by 4.55 times after three years (between T1 and T3). A significant increase of PB was also found in P3, which was 32.6%. There were little changes of plaque burden in the other three patients.

**Fig 3.**
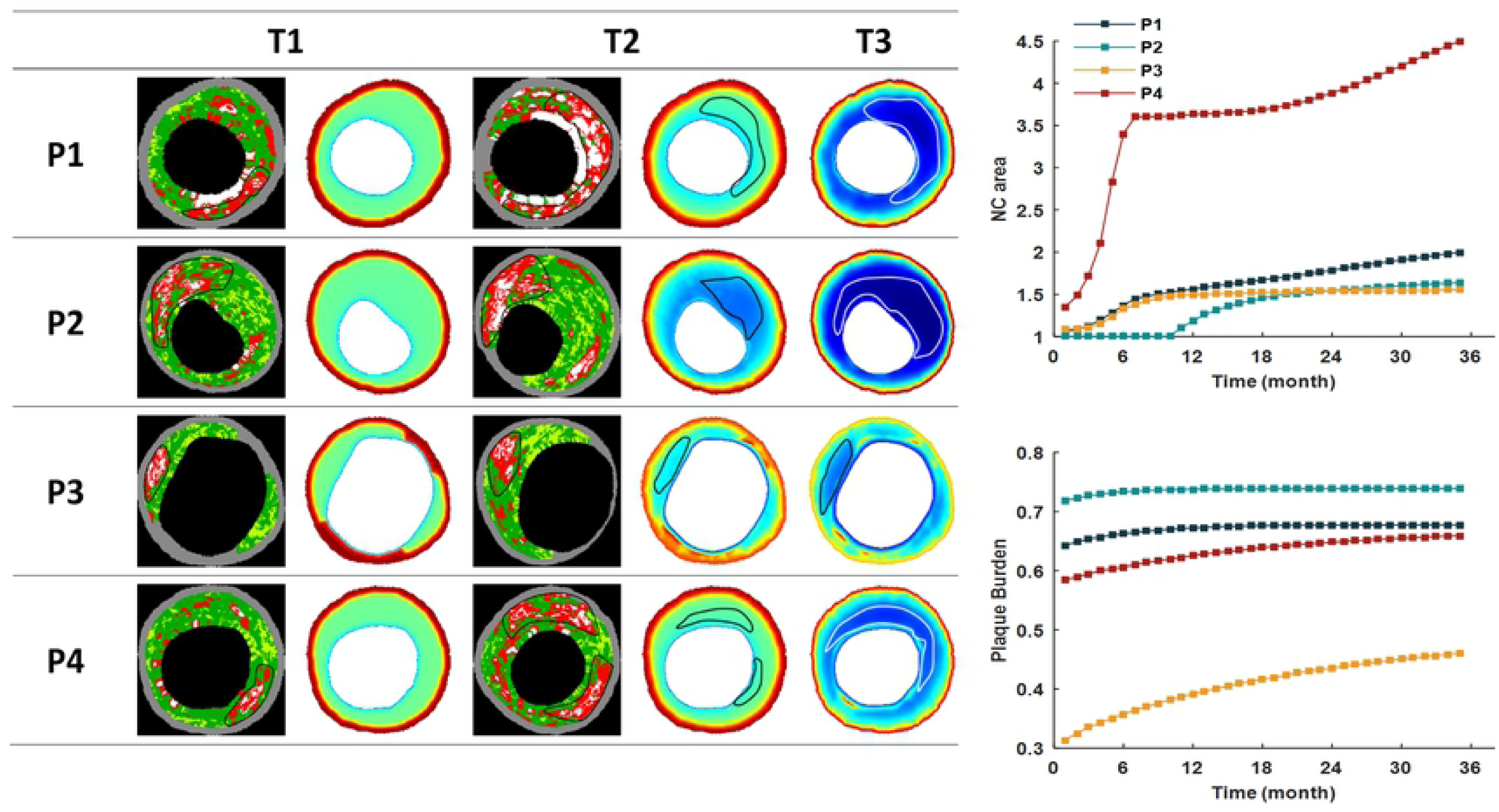
The plaque necrotic core expansion. (A) Four patients’ NC areas were circled in black/white lines in both VH-IVUS images and simulation results. (B) The growth history curves of NC area and PB of the patients for three years. The relative values of NC area were compared with the VH-IVUS data at T1.

Taking into account of systematic and microenvironmental factors, we proposed a scoring scale to predict a possible plaque development (Table 1). The scores of PB and LDL were high in P1, suggesting a severe lipid deposition in the plaque. However, the local hemodynamics was not favorable for plaque development (1 score for wall shear stress, WSS), resulting in a relatively benign microenvironment in P1. In addition, the flaky distributed calcification areas were shown in the VH-IVUS image (white regions). Therefore, a stable plaque with macrocalcification was reckoned as its future growing of P1’s plaque. Although the evaluation of the microenvironmental factors in P3 was quite severe, putting together with the other indictors (plaque morphology and blood test) indicated it was a stable plaque. Consequently, the rate of plaque development in P3 was relatively slow, and the NC area was not remarkable during the plaque progression. Contrary to the predictions of P1 and P3, both P2 and P4 have the highest scores (Total score = 20), which suggested vulnerable plaques were more likely to be found in these patients. In summary, combing with the analysis of VH-IVUS data and the simulation results, P2 and P4 were classified as unstable group, and P1 and P3 as stable group.

### Influence of microenvironmental factors on plaque progression

According to the prediction for plaque progression, P1 and P3 were categorized as the stable group, while P2 and P4 as unstable group. Fig 4 shows the difference of the changes of microenvironmental factors during progression between the two groups. Independent t-test demonstrated that there were significant differences in the five microenvironmental factors (LDL, ox-LDL, MCP-1, SMC, and foam cell, p < 0.01). However, no significant differences were found in monocytes (p = 0.11), macrophages (p = 0.011), and ECM (p = 0.02).

**Fig 4.**
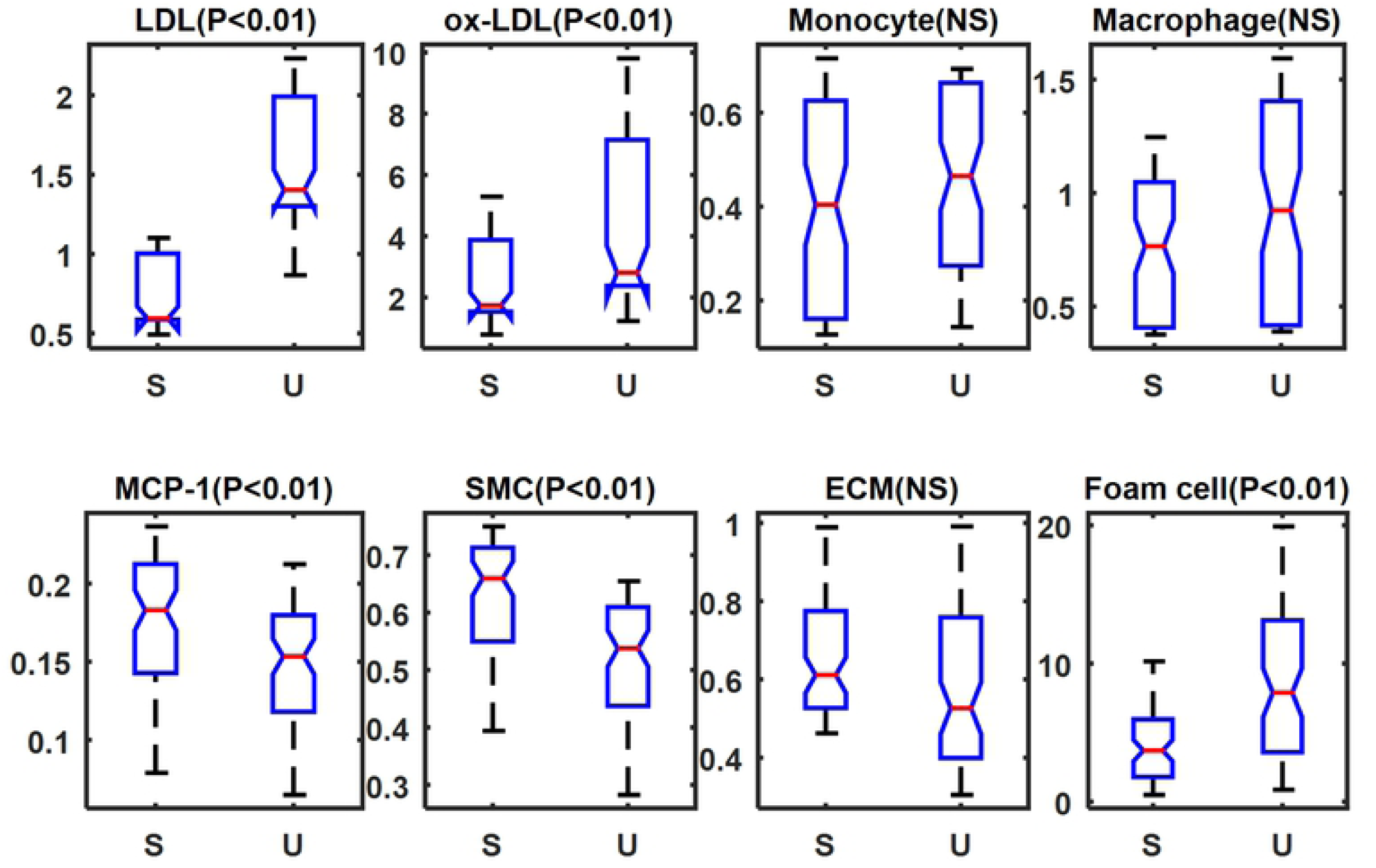
Comparisons of the microenvironmental factors on plaque progression between stable group (S) and unstable group (U). NS = not significant.

Then, we investigated the principal factors that influenced the NC growth of the two groups by using the Spearman’s correlation analysis. As shown in Table 2, in the stable group, LDL, ox-LDL, SMC, and foam cell showed strong corrections with the NC growth (r = 0.7827, 0.8810, -0.8361, 0.8566 respectively). While in the unstable group, monocyte, macrophages, MCP-1, and ECM showed strong correlations with the NC growth (r = 0.9919, 0.8692, 0.7797, -0.8406, respectively). It was noteworthy that the inflammatory microenvironment (monocyte and macrophage) had a significant negative correlation with NC expansion in the stable group (r = -0.3389 and r = -0.3076, respectively). In contrast, it had a very strong positive correlation in the unstable group (r = 0.9919 and r = 0.8692, respectively). This result indicated that the inflammatory factors could be beneficial or detrimental during plaque progression, depending on the interactions with other microenvironmental factors.

## Discussion

Although the formation and progression of atherosclerotic plaques is understood to be mainly driven by the systemic factors, both in vitro and in vivo observations over the past several decades have established that plaque development involves a complex coordination of systemic and local microenvironmental factors that determine how plaques progress. Uncovering the underlying mechanisms and crosstalk of plaque microenvironment constitutes as prerequisite for the exploration of possibilities targeting the processes within the plaque microenvironment as novel therapeutic strategies. Recently, the results of the Canakinumab Anti-inflammatory Thrombosis Outcome Study (CANTOS, targeting the interleukin-1β) trial show conclusive proof that reduction of inflammation by inhibiting the IL-1β pathway activation can significantly lower coronary artery disease morbidity and mortality [17]. However, there is currently a paucity of data on the dynamic interactions of the plaque microenvironmental components, due to the unavailability of imaging techniques to quantitatively investigate the changes of plaque microenvironment in both animal experiments and clinical settings. In this context, mathematical modeling provides a potential way to assess the dynamics of the plaque microenvironment by describing the spatial-temporal changes of key cells and chemicals according to physical principles and known pathophysiological interactions. To this end, we performed the evolutional simulations of plaque microenvironmental dynamics based on patient follow-up data and predicted the development of microenvironmental factors as well as plaque phenotype in patient-specific atheroma on theoretical and quantitative grounds.

With the information of plaque microenvironment as initial input, including plaque morphology and composition obtained by imaging technique and biochemical indicators by blood examination, the dynamics and development of the main microenvironmental factors can be calculated for next three years by using the developed model system. Based on these results, the evolutional progress of a specific plaque can be predicted. The evaluation indicators of plaque microenvironment include lipid deposition, inflammation, IPH and apoptosis of SMCs. The results clearly demonstrate the heterogeneity of the microenvironmental factors within plaque and during the plaque progression. It has been suggested that plaque progression is a modifiable step in the evolution of atherosclerotic plaque [18]. Therefore, a dynamic and quantitative description of plaque microenvironment can provide direct information for personalized treatment to improve the long-term outcomes. In addition, a scoring scale table for each patient is presented, which comprises the dynamic microenvironmental indicators with the classical risk index (morphological and biochemical index). For the very first time, a quantitative evaluation is proposed by a comprehensive consideration of both morphological factors in tissue level and functional factors in cellular level. We believe the improved evaluation system can help us better understand the interactions among plaque microenvironmental factors and may allow us to predict a possible development of a plaque on an individual basis.

According to the prediction for the plaque progression, the patients are divided into two groups, i.e., stable and unstable group. We investigate the differences of the main microenvironmental factors between the two groups. The statistical analysis reveals that the lipid components (LDL and ox-LDL) rather than the inflammatory factors (monocyte and macrophage) exhibit significant differences between the stable plaques and the unstable ones. In addition, the correlation test of the microenvironmental factors with NC enlargement in different groups was performed. In the stable group, the lipid components have a strong positive correlation with the NC expansion, while in the unstable group, the inflammatory factors that have a very strong positive correlation with the NC expansion. This interesting result may be used to explain why not all clinical trials provided a beneficial cardiovascular effect especially in the anti-inflammatory therapy [19]. This also suggests that the patients who would better benefit from these therapies (such as the unstable group in this study) could be identified according to the dynamics of their plaque microenvironment. Although the correlation analyses of these microenvironmental factors with clinical outcomes still need to be validated by more clinical studies, the present results emphasize the heterogeneity of plaque microenvironment between individuals and its complex role in plaque development, and provide a potential path toward the investigation of an improved and targeted atherosclerosis therapy.

Several imaging modalities are currently used in vivo to characterize one or more plaque microenvironmental factors. For instance, dynamic contrast-enhanced MRI (DCE-MRI) is proposed to study the intraplaque microvasculature quantitatively and to test the relationship between adventitial perfusion and IPH, while ^18^F-FDG PET-CT is widely used to evaluate plaque inflammation. Compared with the conventional structural imaging techniques identifying the site and severity of luminal stenosis, these functional assessments may provide more informative values in studying the dynamic microenvironment and consequently to evaluate plaque vulnerability. However, it is unrealistic to carry out multi-modality imaging for every patient due to the technological and socio-economic issues. In this context, this proof-of-concept study aims to present a novel personalized evaluation for coronary atherosclerotic plaque microenvironment by incorporating a generalized mathematical modeling system with patient-specific clinical data. The power of mathematical modeling lies in its ability to reveal the underlying dynamic mechanism and the physical principles that might have been overlooked in previous traditional studies. At its best, mathematical modeling provides quantitative supplementary for the imaging data, and enables us to make predictions and early identify which plaque rupture is likely to occur, and leads to a novel and improved ability to assess plaque vulnerability. This will allow actions to be taken in a timely manner to reduce risk of eventual fatal events on an individual basis. Although we have demonstrated the simulation based on VH-IVUS images in this study, further researches should be addressed by incorporating other available imaging data to expand the application of this mathematical model in clinical assessment.

There are several limitations in this study. First, the predicted plaque progression should be validated by clinical outcomes and more patients’ data should be included to demonstrate the statistical significance of this study. Extensive large-scale patient study will be needed to validate the model before it can be used as a prediction tool in clinic. Second, the current model excluded the calcification that is associated with the apoptosis of macrophages and SMCs, and interacts with other microenvironmental factors such as inflammation. Finally, the balance between pro-atherogenic factors and anti-atherogenic factors in plaque microenvironment was not investigated. Considering the dynamic balance among multiple microenvironmental factors was a major study in itself and its influence on plaque development should be addressed in future work.

In conclusion, an image-based patient-specific multi-physical model is developed which can simulate the spatial-temporal evolution of plaque progression as well as the dynamic variations of plaque microenvironment. This enables us to make predictions and early identify the high-risk rupture-prone plaques, leads to a novel, improved ability to assess plaque vulnerability, and allows actions to be taken in a timely manner to reduce risk of eventual fatal events on an individual basis. It is found that the inflammatory microenvironment has a negative correlation with NC expansion in stable plaques while it has a very strong positive correlation in unstable plaques, suggesting that inflammation may be beneficial or detrimental during plaque progression, depending on the interactions with other microenvironmental factors.

## Methods

### Study design

The proposed simulation system is described in Fig 5, in which the whole process consists of two steps: validation and prediction. Since the initial inflammation and neovascularization in the plaque microenvironment cannot be assessed by IVUS imaging, we first identified the initial microenvironmental factors at baseline (T1) images, by comparing the simulation results calculated from different levels of macrophages concentration and microvascular density with follow-up images at T2. Once the plaque microenvironment for each patient at T1 was determined, the simulation was performed to assess the dynamic changes of the main cellular and acellular components involved in the plaque progression. The prediction was then conducted to obtain the plaque development at the end of simulation, T3 (three years after T1).

**Fig. 1. Schematic diagram for simulation algorithms**.

### Data acquisition

Follow-up IVUS with virtual histology imaging data of coronary plaques were obtained at Cardiovascular Research Foundation (New York, NY) and Washington University at St. Louis, from 4 patients (3M, 1F) who had percutaneous coronary intervention (PCI). The IVUS follow-up was at about 6 months or one year. A synthetic-aperture-array, 20-MHz, 3.2-French catheter (Eagle Eye, In-Vision Gold, Volcano) with motorized catheter pullback (0.5 mm per second) was used to acquire IVUS data. All imaging data were from the PROSPECT study [20] and the research was approved by the Institutional Review Board (IRB) at Cardiovascular Research Foundation and the IRB at Washington University St. Louis, respectively. All patients provided written informed consent.

Level of serum creatinine, fasting lipids, glucose, glycated hemoglobin, and high-sensitivity C-reactive protein were measured at baseline. Fusion of IVUS data and X-ray angiography to reconstruct 3D arterial geometry were performed after the segmentation and co-registration of the one-by-one paired slices at baseline (T1) and follow-up (T2) images. IVUS-based 3D fluid-structure interaction (FSI) models with cyclic bending were constructed for each coronary to assess the flow WSS conditions [21,22]. For each patient, the paired slices with the maximum plaque burden/area were selected in this study. The detailed demographical information of patients was listed in Table 2.

### Image-based patient-specific multi-physical modeling

Based on our recently developed model [15,16], we modeled plaque microenvironment as a continuum medium that the cellular/acellular factors diffuse within the thickening intima as well as interact with each other. In particular, we considered four main pathophysiological processes during plaque development, i.e., lipid deposition, inflammatory response, migration and proliferation of SMCs, and neovascularization. These four pathophysiological phenomena were coupled based on the following experimental and clinical observations: (a) ox-LDL activates the expression of proinflammatory cytokine such as MCP-1 to facilitate the recruitment of more monocytes into the lesion; (b) the neovasculature provides a potential way for LDL and monocytes into the intima by extravasation from the leaky vessel wall; (c) the accumulation of lipoprotein and inflammatory cells (monocytes and macrophages) promotes SMCs migration and proliferation, resulting in a hypoxic microenvironment to induce further angiogenesis in the thickening intima. We modeled the cellular and acellular components involved in the above four processes. Namely, the cellular components consist of endothelial cells (ECs), macrophages, monocytes, foam cells and SMCs, while the acellular counterparts include LDL, ox-LDL, MCP-1, VEGF, MMP, ECM and extravascular plasma concentration. The interactions between all variables in this model are illustrated in Fig 6. The dynamics of these twelve variations satisfy the mass conservation, described by coupled reaction-diffusion equations as follow:

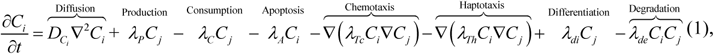

Where *C*_*i*_ denotes one plaque microenvironmental factor, and the first terms on the right-hand side of the equation describes the diffusion of *C*_*i*_ with diffusion coefficient *D*_*Ci*_. The reaction terms of *C*_*j*_ are modeled based on the proved pathophysiological knowledge during the plaque progression, including the production, consumption, chemotaxis, haptotaxis, and differentiation by other microenvironmental Factor *C*_*i*_, as well as the apoptosis by itself. The detailed explanations of reaction terms involved in the model system are listed in Table 3. The numerous parameters involved in the equations of the present model have been estimated from available experimental data and mathematical models wherever possible. For example, *D*_*L*_, the diffusion coefficient of LDL was given by an in vitro experiment that calculated the average radioactivity with time, in the arteries incubated in Tyrode’s solution with radioactive tracer, and fitted the curve in which *D*_*L*_ was a parameter that can be given by the least-squares methods [27]. The coupled equations of all microenvironmental factors in this model are listed in the Supplementary Material. The explanation of the parameter setting and other details can be found in our previous work [15,16]. To avoid the pre-defined interventions as much as possible, we assumed that the interaction coefficients between the microenvironmental factors remain unchanged during plaque progression.

**Fig 6.** The interactions between variables in the model. Seven pathophysiological progresses (circle) among cellular (yellow box) and acellular (green box) factors are included in the model, and every arrow represents a reaction term modelled in the reaction-diffusion equations.

**Table 3.** The detailed explanations of reaction terms involved in the model system

| Variables | Reaction Terms | Explanation |
| --- | --- | --- |
| LDL: $L$ | $-\lambda_L L$ | Lipid oxidation [8] |
| | $\lambda_{Pl_{extra}} Pl_{extra}$ | LDL from leaky microvessels [2] |
| ox-LDL: $L_{ox}$ | $\lambda_{Lox \cdot L} L$ | ox-LDL formed by oxidation of LDL [23] |
| | $-\lambda_{Lox \cdot Ma} L_{ox} Ma$ | Macrophage phagocytosis of ox-LDL [23] |
| MCP-1: $P$ | $\lambda_{P \cdot E} \frac{L_{ox}}{K_P + L_{ox}} E$ | Production of MCP-1 produced by ECs [16] |
| | $\lambda_{P \cdot S} S$ | Production of MCP-1 by SMCs [24] |
| | $-d_p P$ | Decay of MCP-1 [2,24] |
| Macrophage: $Ma$ | $-\nabla(\lambda_{Ma \cdot P} Ma \nabla P)$ | Chemotaxis of macrophages in response to MCP-1 [8] |
| | $\lambda_{Ma \cdot Mo} Mo$ | Macrophages differentiated from monocytes [8] |
| | $-d_{Ma} Ma$ | Apoptosis of macrophages [8] |
| Monocyte: $Mo$ | $\nabla(\lambda_{Mo \cdot L_{ox}} Mo \nabla L_{ox})$ | Chemotaxis of monocytes in response to ox-LDL [8] |
| | $\lambda_{Pl_{extra}} Pl_{extra}$ | Monocytes from leaky microvessels [2] |
| | $-d_{Mo} Mo$ | Decay of monocytes [23] |
| ECs: $E$ | $-\nabla \left( \frac{\lambda_{E \cdot C_v}}{K_E + C_v} E \nabla C_v \right)$ | Chemotaxis of ECs in response to VEGF [2] |
| | $\nabla(\lambda_{E \cdot C_{ECM}} E \nabla C_{ECM})$ | Haptotaxis of ECs in response to the ECM [2] |
| VEGF: $C_v$ | $-\lambda_{C_v \cdot E} E$ | VEGF uptake by ECs [2] |
| | $\lambda_{C_v \cdot S} S$ | VEGF early production by SMCs [2] |
| | $\lambda_{C_v \cdot Ma} Ma$ | VEGF late production by Macrophages [2] |
| | $-d_{C_v} C_v$ | Decay of VEGF [2] |
| Plasma: $Pl_{extra}$ | $-\psi \nabla(U_i Pl_{extra})$ | Convection of plasma [2] |
| | $\gamma Q_t$ | Fluid flux of plasma [2] |
| SMCs: $S$ | $-\nabla(\lambda_{S \cdot P} S \nabla P)$ | Chemotaxis of SMCs in response to MCP-1 [24] |
| | $-\nabla(\lambda_{S \cdot Ma} S \nabla Ma)$ | Chemotaxis of SMCs in response to PDGF secreted from macrophages [24] |
| | $\nabla(\lambda_{S \cdot C_{ECM}} S \nabla C_{ECM})$ | Haptotaxis of SMCs in response to ECM [24] |
| | $-d_S S \cdot L_{ox}$ | Apoptosis of SMCs induced by ox-LDL [25,26] |
| ECM: $C_{ECM}$ | $-\lambda_{C_M \cdot C_{ECM}} C_{ECM} C_M$ | ECM degraded by MMPs [2] |
| | $\lambda_{S \cdot C_{ECM}} S$ | ECM produced by SMCs [24] |
| MMP: $C_M$ | $\lambda_{C_M \cdot E} E$ | MMPs produced by ECs [2] |
| | $\lambda_{C_M \cdot S} S$ | MMPs produced by SMCs [23] |
| | $-d_{C_M} C_M$ | Decay of MMPs [2, 27] |

### Microenvironment analysis

We chose the changes of NC area and plaque burden (area) as the evaluation indicators of plaque development. In the simulation results, the plaque area indicated the area between the lumen and the outer boundary. The NC area was calculated by the apoptotic SMCs, i.e., the area where SMCs decreased more than 50% compared with their initial value.

Since the dynamic distribution of the main factors in the microenvironment can be obtained from the simulation, a scoring scale was proposed to quantify the severity of the atherosclerosis, which comprised the morphological index (plaque burden and eccentric index), the lipid panel test (LDL and HDL), the local hemodynamic factor (WSS), and the representative intraplaque microenvironmental factors (macrophages, SMCs, and ox-LDL). There were three grades on the scale. A larger number indicated a more severe influence on plaque progression (1 score = mild; 2 score = moderate; 3 score = severe). The severity of microenvironmental factors was defined by the changes of each variable during the follow-up interval, while the influence of other risk factors was estimated according to the current consensus. For example, a higher LDL and a lower HDL concentration have been demonstrated as blood indicators of a higher probability that a plaque may develop [7]. And a higher plaque burden and/or eccentric index, as the morphological factors, indicate that the plaque is more dangerous [28–30]. In terms of hemodynamics, it has been found that low and/or oscillatory shear stress contributes to atherogenesis.

## Acknowledgement

This research is supported by the National Nature Science Foundation of China (Grant numbers 11972118, 61821002, 11772093, 11972117), the Fundamental Research Funds for the Central Universities, the National Demonstration Center for Experimental Biomedical Engineering Education (Southeast University, Nanjing, China), the Funds for Young Zhishan Scholars (Southeast University, Nanjing, China), and ARC (Grant number FT140101152, DP200103492)

## Supporting information

**S1 File. Supplementary methods and results**.

